# Bitter taste perception in BaYaka hunter-gatherers

**DOI:** 10.1101/2021.11.25.469973

**Authors:** Sarai Keestra, Edmond Sylvestre Miabangana, Nikhil Chaudhary, Inez Derkx, Gaurav Sikka, Gul Deniz Salali

**Affiliations:** Department of Epidemiology and Data Science, Amsterdam UMC, University of Amsterdam, Amsterdam, the Netherlands; Paediatric Endocrinology, Emma Kinderziekenhuis, Amsterdam UMC, University of Amsterdam, Amsterdam, the Netherlands; Agence Nationale de Valorisation des Résultats de la Recherche et de l’innovation (ANVRI), Cité de la Science de Brazzaville, Brazzaville, Republic of Congo; Leverhulme Centre for Human Evolutionary Studies, Department of Archaeology, University of Cambridge, Cambridge, United Kingdom; Department of Anthropology, University of Zürich, Zürich, Switzerland; NHS Greater London, London, United Kingdom; Department of Anthropology, University College London, United Kingdom

**Keywords:** traditional medicine, plant ecology, dietary transition, PTC, thiourea, hunter-gatherers, bitter taste

## Abstract

Aversion towards bitter tastes evolved across vertebrate species to enable the recognition of harmful plant toxins. Most studies to date have investigated the variation in bitter taste sensitivity between human populations. However, there is a lack of research investigating phenotypic plasticity and the variation in bitter taste perception within the same population. Here we examined bitter taste perception among the Mbendjele BaYaka hunter-gatherers from Congo, a group of forest hunter-gatherers who exhibit a variation in their levels of market integration. We conducted an experiment using phenylthiocarbamide (PTC) and thiourea infused paper strips and compared the prevalence of bitter tasting phenotypes between the BaYaka who grew up in town and forest camps. We found that 45.1% of BaYaka experience PTC as bitter, and 42.5% experience thiourea as bitter. There were no sex and age differences in bitter taste perception. Despite a shared genetic background, we found that BaYaka who grew up in town were more sensitive to bitter taste than those living in the forest, suggesting a developmental component in taste perception. We suggest that a decreased use of traditional plant medicine in town-born BaYaka may underlie this variation in bitter taste perception.

## Introduction

Bitter taste perception evolved across vertebrate species to detect the presence of potentially toxic molecules in plants and evoke avoidance behaviour from such materials [1–3]. Aversion to bitter tastes is innate [4], but bitter detection threshold may increase with age [5]. Bitter taste perception is enabled by ∼25 genes of the TAS2R family of G-protein-coupled receptors that are expressed on the microvilli of taste bud cells on the tongue [6,7]. TAS2Rs help recognise a range of plant compounds including, but not limited to, alkaloids [8,9], saponins [10], tannins [11], flavonoids [12], flavonols [13], glucosides [14], and thioamides [1]. Some compounds have a negative effect on health, for example goitrogenic thioamides, such as thiourea and goitrin [1], and may cause the enlargement of the thyroid gland, and increase thyroid cancer risk [15,16]. It is therefore evolutionarily advantageous to detect such bitter compounds to avoid their ingestion, and nearly all naturally occurring plant poisons are perceived by humans as bitter [11,17]. However, not all bitter tasting foods contain such poisons, and there are also many bitter tasting compounds that are harmless when consumed in the right doses. For example, some naturally occurring bitter compounds such as caffeine in coffee, polyphenols in green tea, and flavonoid branched-chain glycosides in citrus fruits taste bitter but are not harmful to human health in small quantities [7,13,18,19]. Additionally, many medicinal compounds, such as willow bark derived acetylsalicylic acid, the active compound in aspirin, taste bitter and activate the TAS2R bitter receptors, but are beneficial for health when consumed in small doses [7,20,21]. It is therefore the omnivore’s dilemma to discriminate effectively between toxic bitter foods and those that are harmless or even beneficial [14,22].

In the 1930s it was discovered that variation existed in the way humans perceived the taste of the synthetic compound phenylthiocarbamide (PTC) [23]. PTC binds to the same bitter taste receptor as the thioamides [1]. PTC infused paper strips have since been used to test phenotypic ability to perceive bitter tasting compounds. Population variation in PTC sensitivity is found ranging from 68.5% of white Americans of European descent perceiving PTC as bitter [24], to 93.9% of Native Americans [25]. This has brought particular attention to the role of diet and subsistence strategy in understanding this variation [26]. Throughout the history of domestication humans selected against toxic and distasteful plants, and as a result, cultivated plants that are less bitter than their wild counterparts [27,28]. Dietary changes in the bitterness of foods consumed may have in turn affected the strength of selection of bitter taste loci in some agricultural populations, while a diet rich in plant toxins may select for increased bitter taste sensitivity [14]. Sjöstrand *et al*. (2020) looked at bitter taste perception among Baka hunter-gatherers and neighbouring Bantu farmers in Cameroon, comparing their ability to detect bark-derived bitter quinine, which is used as a malaria treatment and activates TAS2Rs [29,30]. Although one may have predicted that a population practising hunting and gathering in a complex forest ecology would benefit from increased sensitivity to plant toxins, they found that Baka were less sensitive to this bitter substance than the neighbouring farmers. However, whether subsistence strategy accounts for these differences remains debated. Campbell *et al*. 2017 investigated variation in bitter taste perception across populations in West, Central, and East Africa with diverse subsistence strategies, ranging from hunting–gathering and pastoralism to agriculture [31]. Although there was a large variation in bitter taste perception across these populations, TAS2R38 haplotype variation, which accounts for 55-85% of the phenotypic variation in responses to bitter compounds [32], was not found to be correlated to geography or diet [31].

One explanation for the contradictory results regarding the association between bitter taste variation with geography or diet may be because bitter taste perception is a more plastic trait than assumed in these studies. Although it is known from studies in rodents and humans that repeated exposure to bitter tastes over several days or weeks can increase acceptance [33–36], no study thus far has investigated the variation of bitter taste perception within a single population where the consumption and use of wild plants vary. The possibility that differences in wild plant use during development within the same genetic population may affect bitter tasting phenotypes remains to be explored. To address this gap, we examined whether a population of forest hunter-gatherers exhibit variation in bitter taste perception based on the frequency of their wild plant use. We conducted a bitter taste perception experiment amongst the Mbendjele hunter-gatherers, who are a subgroup of BaYaka hunter-gatherers living in the rainforests of the Republic of Congo. Consumption of wild plants for medicinal use is traditionally very important amongst the BaYaka [37,38], who are known for their rich knowledge and diverse exploitation of wild plants [38].

The BaYaka often use bitter tree barks for treatment purposes, boiling them and consuming the juice for digestive disorders (mostly caused by parasitic infections), one of the most common illnesses among forest hunter-gatherers [37]. The BaYaka exhibit variation in the levels of market integration and subsistence strategies – in recent decades some BaYaka are born and raised in large settlements near farmer villages and market towns and alongside foraging, engage in gardening and wage labour, while other BaYaka grow up in smaller camps in the forest, relying more heavily on hunting and gathering [39]. Those who grew up in town also report knowing and using fewer wild plants and preferring western medicine over traditional plant medicine [40]. Considering the frequent consumption of such bitter plants in the forest and the previous findings by Sjöstrand *et al*. 2020 in another Congo Basin hunter-gatherer population [29,38,40], we hypothesised that the BaYaka may be less sensitive to bitter taste if they more frequently consume wild plants and more regularly make use of traditional plant medicine. By conducting a bitter taste experiment in town-born versus forest-born BaYaka hunter-gatherers, we tested if exposure to bitter substances during development affects variation in bitter taste perception.

## Methods

### Fieldwork and study population

We conducted fieldwork in three different BaYaka campsites in the northern Republic of Congo in July and August 2018. One campsite was located near a logging town, Pokola, and consisted of individuals who are born and raised there, and families who temporarily move there to visit relatives or buy products in the market. Here we exclusively recruited individuals that were born and grew up in Pokola. The BaYaka who are born and raised in town engage in wage labour more frequently [39], know fewer wild plants [27], and prefer western medicine over traditional treatment practices [40] compared to the forest-born BaYaka. We also recruited individuals from two forest camps (Longa and Njoki), located in the forest near a dirt road. The BaYaka living in forest camps primarily engage in hunting and gathering, but also trade some forest items with Bantu farmers in return for cigarettes, alcohol, and cultivated foods. Nevertheless, BaYaka living in the town can more easily access cultivated foods through Bantu shops and the local market in Pokola.

### Taste perception experiment

We used paper strips containing either phenylthiocarbamide (PTC) or thiourea compounds, in a concentration of approximately ∼30 mcg/strip, and plain papers (as a control), produced by Eisco Laboratories. Participants were given a control paper first, then PTC and finally thiourea. Participants were asked to describe the taste of the papers in an open-ended question. Only individuals that experienced the control paper as tasteless, sweet, or salty were included in the analyses, whereas those who described the control as bitter were excluded. Interviews were conducted in Mbendjele (the language spoken by the study population) with the help of a Mbendjele translator who translated from French to Mbendjele.

### Study sample

Of the 143 participants, 30 did not pass the control test. All other participants, 58 women and 55 men, living in the forest camps (n=74) or the town camp (n=39) were included in the study.

### Data analysis

Mean percentages of tasters of PTC and thiourea were calculated, where “tasting” was defined as individuals that described experiencing the PTC or thiourea paper using terms such as ‘bitter’, ‘a bit bitter’, or ‘very bitter’ in their answer. “Non-tasting” was defined as people describing the taste as nothing, sweet, or salty. We performed logistic regression analyses to examine the effect of camp residence, age, and sex on the odds of PTC and thiourea bitter tasting (tasters coded as 1, and non-tasters as 0). Analyses were conducted in R Studio version 1.3.959.

## Results

### Less than 50% of the BaYaka has a bitter tasting phenotype

Of the 113 individuals included, 45.1% (51/113) experienced the PTC paper as bitter, 50.4% (57/113) were non-tasters of PTC, describing the PTC paper as being tasteless, sweet, or salty, and a small proportion (4.4%; 5/113) described it as hot (Supplementary 1). Additionally, 42.5% (n=48/113) experienced the thiourea paper as bitter, 55.7% (63/113) as nothing, sweet, or salty, and 1.8% (2/113) as hot. There were 26 individuals (23%) who described both PTC and thiourea as bitter. Nevertheless, thiourea tasting did not predict PTC tasting (Supplementary Table 1).

### Bitter tasting phenotype is more frequent amongst town-born hunter-gatherers

Because there was no significant difference between the proportion of tasters in the two forest camps, we combined their responses (PTC: Longa: n=16/49 and Njoki:10/25; thiourea Longa 15/49 and Njoki = 8/25). The percentage of bitter tasters for both the PTC and thiourea was higher among town-born BaYaka than the participants born in forest camps (Figure 1). Our regression models showed that town-born BaYaka were three times more likely to taste PTC and thiourea as bitter than BaYaka who were born in the forest (Table 1).

**Figure 1.**
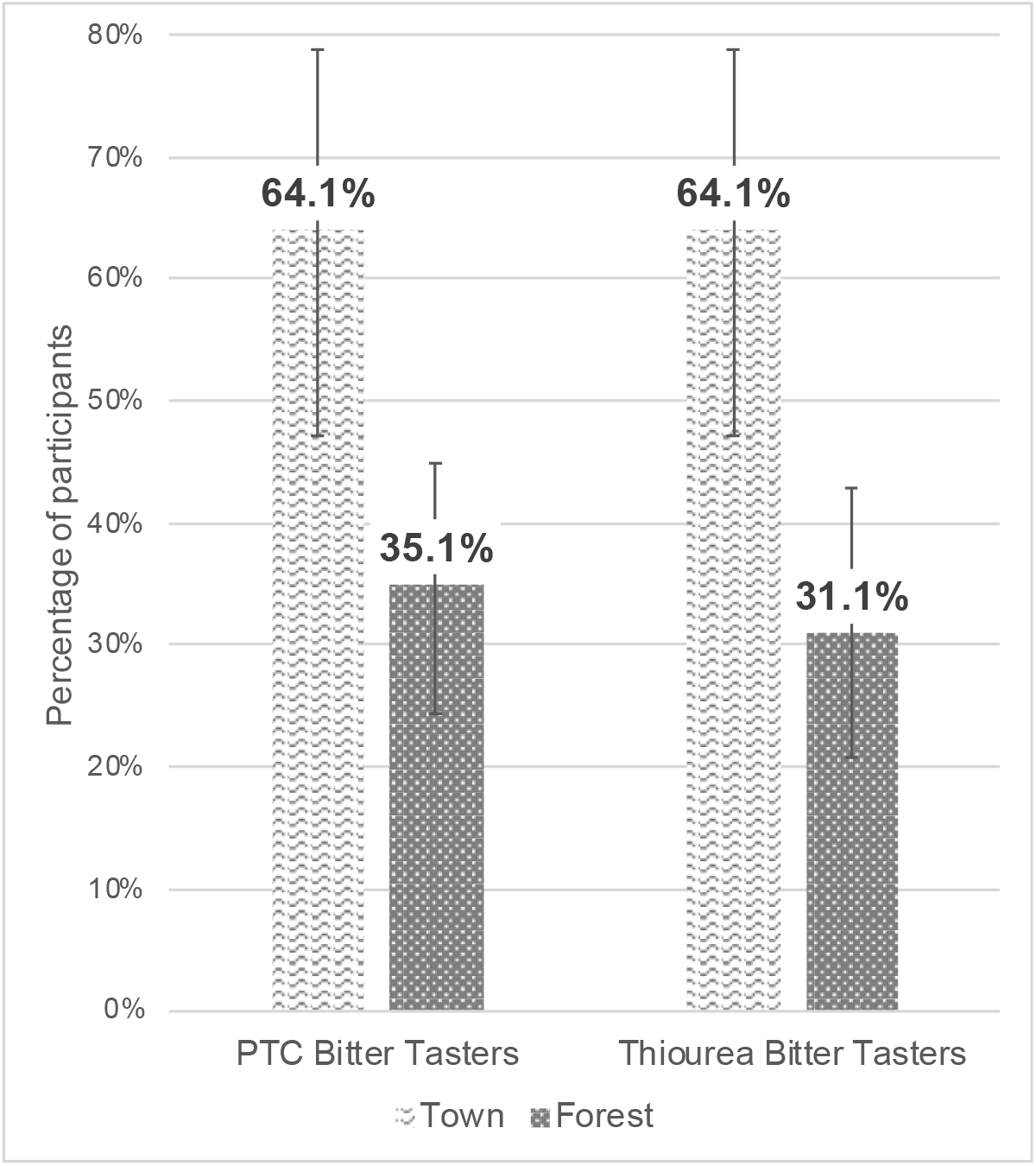
Percentage of PTC and thiourea bitter tasters in BaYaka hunter-gatherers by birthplace (town-born BaYaka n= 39, forest-born BaYaka n= 74).

**Table 1.** Logistic regression models for PTC and Thiourea bitter tasting in BaYaka hunter-gatherers by birthplace and sex.

|  | Odds ratio | 95% Confidence Interval | P-value |
| --- | --- | --- | --- |
| **PTC** |  |  |  |
| Town vs forest | 3.04 | 1.35-7.09 | **0.008** |
| Men vs women | 0.56 | 0.25-1.23 | 0.151 |
| **Thiourea** |  |  |  |
| Town vs forest | 3.85 | 1.71-9.00 | **0.001** |
| Men vs women | 0.68 | 0.30-1.50 | 0.342 |

### No sex and age differences in bitter taste perception amongst the BaYaka

There were no significant differences in PTC and thiourea tasting according to sex (Figure 2, Table 1). When we conducted an additional analysis with the subset of the data where we had age estimates of participants, we found no association between age and bitter tasting (Supplementary Tables 2 and 3).

**Figure 2.**
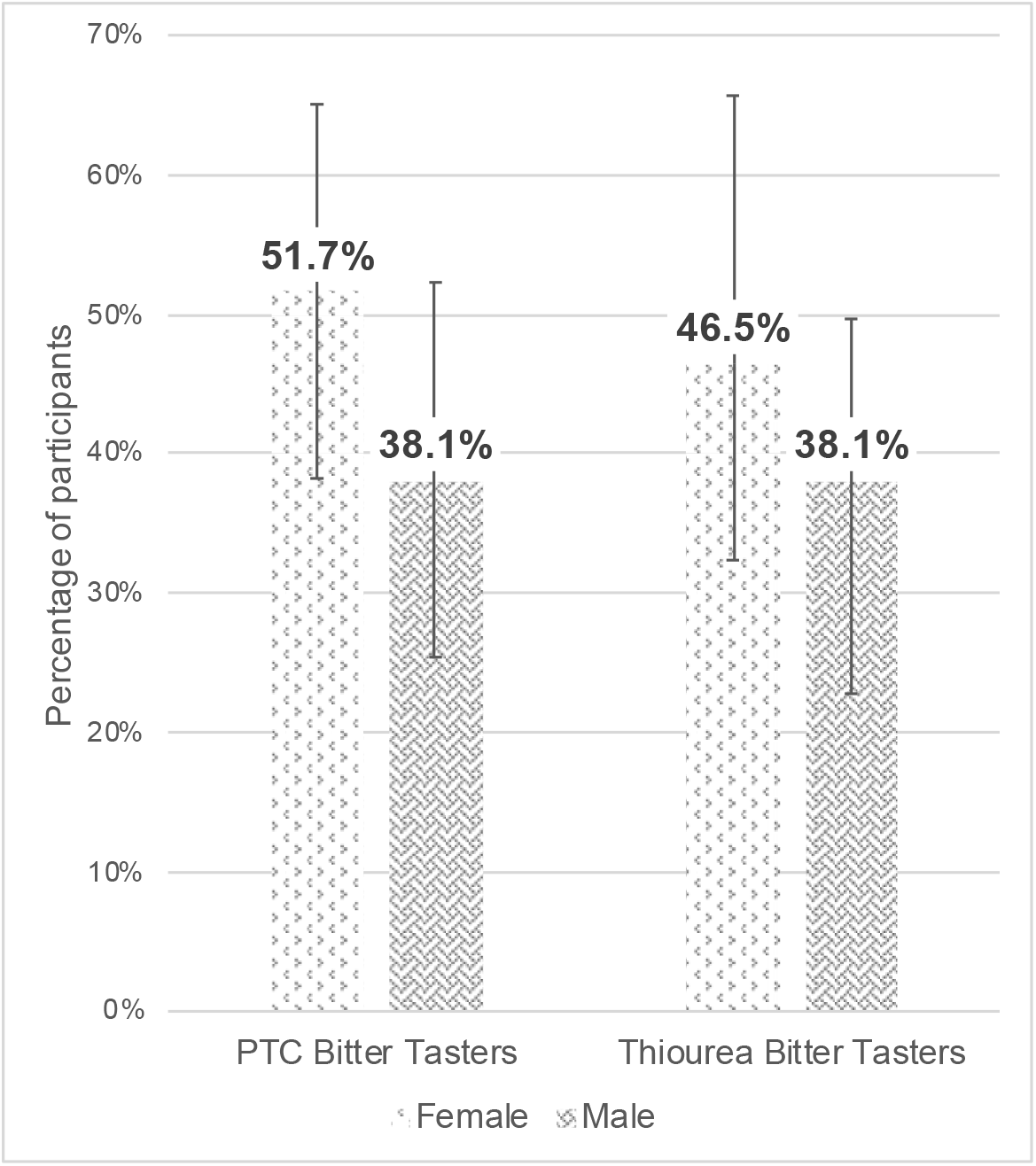
Percentage of PTC and thiourea tasters in BaYaka hunter-gatherers by sex (58 females, 55 males).

## Discussion

In this study, we investigated bitter taste perception among the BaYaka hunter-gatherers. Our results show that a higher percentage of BaYaka hunter-gatherers who were born and raised in town than the forest-born BaYaka perceived the PTC and thiourea as bitter. The PTC bitter tasting phenotype prevalence of 64% in town is more similar to the prevalence of the bitter tasting phenotype found in Western populations of European ancestry (∼60-70%) [16,29], whereas in forest camps 35% were PTC bitter tasters. For thiourea the tasting prevalence was even lower, as only 31% of BaYaka who grew up in the forest camps reported this substance as tasting bitter. Our findings suggest that sensitivity to bitterness may vary in adulthood, despite a shared genetic background, according to the chemical ecology encountered during development.

Both tasting and non-tasting phenotypes might have evolutionary advantages [42,43]. For example, non-tasters may be less fussy and more open to exploring a variety of different plant-based foods [21,32], or better able to ingest a wider range of plant compounds [42,43]. A possible explanation for differences in bitter taste perception between forest and town-born BaYaka could be that although the BaYaka in the forest may consume a diet that is higher in various bitter plant compounds, it is possible to learn to tolerate particular types of bitter compounds if one is exposed to them during early life and if they confer benefits when consumed in small quantities. In non-human primates, for instance, decreased sensitivity to bitter tastes has evolved as an adaptation to leaf-eating behaviour [1,45]. The BaYaka use and consume plants, especially bitter barks, mostly for medicinal purposes [37,38]. In communities where plant treatments are commonly used as medicine, from the ancient Greeks to the agro-pastoralist Arbëresh peoples, the potency of medicinal plants is often derived from the bitterness of their taste [26,46]. Indeed, when asked, the Arbëresh distinguish between plants meant for consumption and those meant for medicinal use by classifying them by their degree of bitterness [46]. Compared to those born in town, BaYaka living in the forest use more wild plants for medicine and prefer traditional treatment methods which often involve boiling of barks and consuming the bitter juice [37,38,40]. It has been previously documented that there is an overlap between the plant species used by the BaYaka and Baka hunter-gatherers and plants used by gorillas and chimpanzees inhabiting the same region for the purpose of self-medication [38]. Previous research has furthermore highlighted a connection between the ingestion of bitter plants among chimpanzees and their anti-parasitic effects [14,47].

To further examine the bitter tasting compounds in the wild plants that are frequently used by the forest-born BaYaka, but not town-born BaYaka, we conducted a literature review. Supplementary Table 4 contains those plants, their frequency of reported usage by forest versus town-born BaYaka (using the data collected by the senior author in 2014 and reported in [40]), their use purpose and the presence of bitter tasting compounds. Our review highlighted that each of those medicinal plants contained bitter compounds such as alkaloids, saponins, and tannins. Most of those plants also have documented bioactive properties and the majority of the medicinal uses concern treating digestive and respiratory disorders and infections [38]. Therefore, in the high-pathogen ecology of the BaYaka living in the forest, it may be advantageous to be less aversive to the bitter taste of plants and barks containing medicinal properties that may confer a survival advantage when consumed at low doses. This aligns with the findings of Sjöstrand *et al*. (2020) who found the Baka in Cameroon to be less sensitive to bark-derived quinine than neighbouring farming groups [29]. Based on the findings of the taste experiment presented here, we hypothesise that as a result of the chemical ecology that BaYaka encounter as children, the tolerance and therefore detection threshold for bitter tastes amongst those who grew up in the forest may be higher.

Our taste experiment suffered from several limitations. Firstly, our sample size was small as the BaYaka mostly lived in groups fewer than 60 individuals [48]. Secondly, we did not exclude individuals that described the paper as sweet or salty at the control stage as this may reflect the flavour of plain paper to an unaccustomed taster. Nevertheless, we found that more stringent exclusion criteria did not affect the significance of associations found between bitter taste perception and birthplace (Supplementary Table 5). Thirdly, we only used a single concentration of PTC and thiourea, which precluded an assessment of whether there are differences in bitter taste threshold between the forest and town BaYaka groups. Additionally, further research is needed to verify whether the bitter taste responses found in response to two artificial compounds translate to the perception of other bitter foods and medicines BaYaka encounter in their everyday life and environment. As most BaYaka do not know their chronological age based on the Western calendar, this limited our ability to look at the effect of age on bitter taste perception in this population. Finally, anthropological studies on the terminology of different tastes and gradations of “bitter” in the BaYaka are needed to better understand the gradient of perception of bitter compounds in this population and how it is correlated with food and treatment choices.

## Conclusion

In a hunter-gatherer population, we found that individuals living in the forest are less sensitive to bitter taste than those who grew up in a town with reduced use of medicinal plants. Our findings challenge the commonly held assumption that bitter taste perception is genetically determined and suggests the existence of a developmental component in bitter tasting phenotype.

## Acknowledgements

We would like to express our thanks to all Mbendjele BaYaka who participated in our study. We are grateful for their participation and hospitality during the project. Our particular thanks go to our translators Nicolas, Roger, and Mindoula for their invaluable help in data collection. We also would like to thank Jerome Lewis, Clobite Bouka Biona, and Laure Stella Ghoma Linguissi for their help with fieldwork logistics and research permits.

## Author contributions

G.D.S conceived the project; S.K, G.D.S., N.C., G.S., I.D., collected the data; E.S.M. helped with the literature review on bitter compounds found in medicinal plants; S.K. conducted the analyses and wrote the manuscript under the supervision of G.D.S.; all authors reviewed and approved the final manuscript.

## Data availability

Data associated with this study is available at OSF: https://osf.io/6t4vd/

## Ethics

The research and fieldwork were approved by the Ethics Committee of University College London (ethics code: 13121/001) and the methods were carried out in accordance with the approved guidelines. Informed consent was obtained from all participants and research permission was granted by the Republic of Congo’s Ministry of Scientific Research.

