## Supplementary Table 1 for "Bitter taste perception in BaYaka hunter-gatherers"

These are the results for the analysis of bitter taste perception in BaYaka hunter-gatherers examining the overlap between PTC and thiourea bitter tasting. We did not find that being able to perceive one compound correlated significantly with tasting the other (p=0.08; OR: 2.00; 95% confidence interval: 0.93-4.41). We recount here who of the bitter tasters thought that the other compound was also bitter. Of the PTC bitter tasters, 26 found thiourea also bitter, 24 did not and 1 found it hot. Of the thiourea bitter tasters, 26 found PTC also bitter, 20 did not and 2 found it hot. To conclude, there were 26 participants who experienced both compounds as bitter. In the regression below, we examined whether thiourea tasting predicted PTC tasting controlling for sex and birthplace. The only significant variable was the birthplace; perceiving thiourea as bitter did not predict perceiving PTC bitter (Supplementary Table 1).

Supplementary Table 1. Odds of PTC tasting with respect to thiourea tasting, sex and birthplace

|  | Odds ratio | 95% Confidence Interval | P-value |
| --- | --- | --- | --- |
| Thiourea tasting | 1.44 | 0.61-3.33 | 0.40 |
| Men vs women | 0.60 | 0.26-1.33 | 0.23 |
| Town vs forest | 2.80 | 1.19-6.80 | 0.02 |

**Age is not significantly associated with bitter taste perception**

Below are the results for the analysis of bitter taste perception in BaYaka hunter-gatherers when age is included in the regression model. Note that not for all participants age was available, as BaYaka do not record their age in years such as in the Western cultural tradition. We had age estimates for 44 participants from the forest camps. Those age estimates were calculated based on the method described in Diekmann et al 2017*.

We conducted a regression analysis with this subset of the data where we had age estimates. We did not find a significant association between age and bitter taste perception in this population (Supplementary Table 2).

Supplementary Table 2. Odds of PTC and Thiourea tasting with respect to age

|  | Odds ratio | 95% Confidence Interval | P-value |
| --- | --- | --- | --- |
| PTC | | |  |
| Age (years) | 0.99 | 0.95-1.03 | 0.71 |
| Thiourea | | |  |
| Age (years) | 0.97 | 0.93-1.01 | 0.16 |

Next, we ran regression models including both age and sex. The results are shown in the table below. There was no association of age and bitter taste perception, controlling for sex.

Supplementary Table 3. Odds of PTC and Thiourea tasting with respect to age and sex

|  | Odds ratio | 95% Confidence Interval | P-value |
| --- | --- | --- | --- |
| PTC | | |  |
| Sex | 0.76 | 0.20-2.82 | 0.68 |
| Age | 0.99 | 0.95-1.03 | 0.71 |
| Thiourea | | |  |
| Sex | 0.33 | 0.08-1.16 | 0.09 |
| Age | 0.97 | 0.93-1.01 | 0.16 |

*Diekmann, Y., Smith, D., Gerbault, P., Dyble, M., Page, A. E., Chaudhary, N., Migliano, A. B., et al. (2017). Accurate age estimation in small-scale societies. *Proceedings of the National Academy of Sciences of the United States of America*, *114*(31).

**Bitter tasting compounds in plants frequently used by the forest born but not town born BaYaka**

Supplementary Table 4. Plants that are not used as frequently by the BaYaka who were born in the town and their bitter tasting compounds

| Plant (BaYaka name) | Latin name | Common Usages | Bitter compounds | Percentage of people using plant as medicine born in forest | Percentage of people using plant as medicine born in town |
| --- | --- | --- | --- | --- | --- |
| Bulaki | [*Caloncoba welwitschi*i Oliv.](https://www.ville-ge.ch/musinfo/bd/cjb/africa/details.php?langue=fr&id=16721) | Headache, pain and other injuries | The bitter taste of the plant is due to the presence in the seeds of chaulmoogric acid, formerly used in the treatment of leprosy [1]. Additionally, the presence of hydrocyanic acid has been detected [2]. | 63% | 14% |
| Indengo | *Croton haumaniamus* J. Léonard (Euphorbiaceae) | Respiratory problems | *Croton* trees are high in active alkaloids [5]. The bitter bark of the related *Croton eluteria* is used in the traditional medicine of northern South America as well [5]. | 93% | 28% |
| Jongo | *Ricinodendron heudelotii* (Baill.) Pierre ex Heckel (Euphorbiaceae) | Infections, pain and injuries | The seeds from the fruit of the *Ricinodendron* have a spicy/peppery taste[6]. | 54% | 10% |
| Kombo | *Musanga cecropioides* (Urticaceae) | Respiratory and digestion problems. | Extract of this nettle type plant contains alkaloids terpenes and saponins which have a bitter and acrid taste [7]. There is a presence of saponins and tannins in trunk and root bark [2]. | 60% | 34% |
| Moba | [*Pentaclethra macrophylla* Benth.](https://www.ville-ge.ch/musinfo/bd/cjb/africa/details.php?langue=fr&id=70031) (Fabaceae) | Digestive problems. In the regions of Brazzaville and Kinshasa, the aqueous decoction of the bark of the trunk is used in the treatment of lumbar and intercostal pain. | The stem bark contains high concentrations of bitter saponins, tanins, and alkaloids [8,9]. The root contains saponins, numerous polyphenols (tannins) and steroids [10]. | 67% | 41% |
| Mongo | *Zanthoxylum tessmannii*  See under:  [*Zanthoxylum gilletii* (De Wild.) P.G. Waterman](https://www.ville-ge.ch/musinfo/bd/cjb/africa/details.php?langue=fr&id=89769) (Rutaceae) | Digestion. Mix the grated bark with cold water and drink the juice. | Bitter tasting alkaloidal (peroxysimulenoline, sanguinarine, xanthoplanine, fagarine I and norchelerythrine), monoterpenes (myrcene, limonene, and camphene) compounds have been isolated from *Zanthoxylum* species [11]. The phytochemical study of a coumarin extract of Zanthoxylum gilletii leaves revealed in addition to coumarins, the presence of flavonoids and anthracene derivatives [12]. | 71% | 28% |
| Mokula | *Microdesmis puberula*  *(Pandaceae)* | Erectile dysfunction. Eat the root. | Alkaloid traces found in stems and roots of this plant [13,14], which are typically perceived as bitter [15]. | 61% | 17% |

15.	Nissim I, Dagan-Wiener A, Niv MY. In press. The Taste of Toxicity: A Quantitative Analysis of Bitter and Toxic Molecules. (doi:10.1002/iub.1694)

**Exclusion of participants tasting the control as sweet does not change results**

Below are the results for the analysis of bitter taste perception in BaYaka hunter-gatherers excluding the participants who taste the control paper as sweet or salty. Being more stringent in the exclusion criteria did not change the results or the significance of the associations found.

Of the 113 participants, 27 described the paper as being a bit or very sweet. For the analyses below these participants were excluded, leaving a final sample size of 41 women and 45 men. In total we included 59 individuals from forest camps (38 from Longa and 21 from Njoki) and 27 from the town camp in Pokola.

### Of the 86 individuals included, 45.3% (39/86) experienced the PTC paper as bitter, 52.3% (45/86) were non-tasters of PTC, describing the PTC paper as being tasteless, sweet, or salty, and a small proportion (2.3%; 2/86) described it as hot (Supplementary 1). Additionally, 40.1% (35/86) experienced the thiourea paper as bitter, 58.1% (50/86) as nothing, sweet, or salty, and 1.7% (1/86) as hot.

### *Bitter tasting phenotype is more frequent amongst town-born hunter-gatherers*

The percentage of bitter tasters for both the PTC and thiourea was higher in town-born BaYaka than the participants from the forest camps. Indeed amongst town-born BaYaka 70.4% (19/27) experienced PTC as bitter, compared to 33.9% (20/59) amongst forest-born BaYaka. Similarly, 63.0% (17/27) town-born BaYaka descrived thiourea as bitter, compared to 30.5% (18/59) forest-born BaYaka. Our regression models showed that town-born BaYaka were three times more likely to taste PTC and thiourea as bitter than BaYaka living in the forest (Supplementary Table 5). There was no significant difference in PTC and thiourea tasting according to sex (Supplementary Table 5).

Supplementary Table 5: Logistic regression modes for PTC and Thiourea bitter tasting by camp location and sex with the subset of data.

|  | Odds ratio | 95% Confidence Interval | P-value |
| --- | --- | --- | --- |
| PTC | | |  |
| Town vs forest | 4.41 | 1.67-12.55 | <0.01 |
| Men vs women | 0.56 | 0.22-1.39 | 0.22 |
| Thiourea | | |  |
| Town vs forest | 3.75 | 1.46-10.1 | <0.01 |
| Men vs women | 0.84 | 0.34-2.09 | 0.71 |
